# Behavior-specific occupancy patterns of Pinyon Jays (*Gymnorhinus cyanocephalus*) in three Great Basin study areas and significance for pinyon-juniper woodland management

**DOI:** 10.1101/2020.07.31.230367

**Authors:** John D. Boone, Chris Witt, Elisabeth M. Ammon

## Abstract

The Pinyon Jay is a highly-social, year-round inhabitant of pinyon-juniper woodlands in the western United States. Range-wide, Pinyon Jays have declined ~ 3 – 4% per year for at least the last half-century. At the same time, large acreages of pinyon-juniper woodland have been removed or thinned to improve habitat for Greater Sage-Grouse or other game species across much of the Great Basin, which is home to nearly half of the global population of Pinyon Jays. Occupancy patterns and habitat use of Pinyon Jays have not been well characterized across much of the species’ range, and obtaining this information is necessary for better understanding the causes of ongoing declines and determining useful conservation strategies. Our goal of this study was to identify the characteristics of areas used by Pinyon Jays for several critical life history components and to thereby facilitate the inclusion of Pinyon Jay conservation measures in the design of vegetation management projects. To accomplish this, we studied Pinyon Jays in three widely separated study areas using radio telemetry and direct observation, and measured key attributes of their locations and a separate set of randomly-selected control sites using the U. S. Forest Service’s Forest Inventory Analysis protocol. Data visualizations, non-metric dimension scaling ordinations, and logistic regressions of the resulting data indicated that Pinyon Jay occupancy was concentrated in a distinct subset of available pinyon-juniper woodland habitat, and further that Pinyon Jays used different habitats, arrayed along elevational and tree-cover gradients, for seed caching, foraging, and nesting. Caching was concentrated in low-elevation, relatively flat areas with low tree cover; foraging occurred at slightly higher elevations with moderate tree cover, and nesting was concentrated in somewhat higher areas with greater tree cover and higher stand density. All three of these Pinyon Jay behavior types were highly concentrated within the lower-elevation band of pinyon-juniper woodland close to the woodland-shrubland ecotone. Because woodland removal projects in the Great Basin are often concentrated in these same areas, it is critical to incorporate conservation measures informed by Pinyon Jay occupancy patterns into existing woodland management paradigms, protocols, and practices.

## Introduction

The Pinyon Jay (*Gymnorhinus cyanocephalus*) is a highly-social corvid that inhabits pinyon-juniper and other coniferous woodlands in the interior western U. S. [1–3] (Fig 1). Pinyon Jays form year-round flocks that can range from a few dozens to several hundred members [3–6]. They are perhaps best known for harvesting and caching the seeds, or “pine nuts”, of the pinyon pine (primarily *Pinus monophylla* and *P. edulis*) as their primary food source, though they also consume other conifer seeds and insects [3, 7, 8]. Pinyon Jays are present year-round in parts of at least ten states, but most of their range lies within Bird Conservation Regions (BCRs) 9 (“Great Basin”) and 16 (“Southern Rockies and California Plateau”) [9], with highest densities in central-eastern Nevada and western New Mexico (Fig 2). Since North American Breeding Bird Survey data collection began in 1967, Pinyon Jays have experienced steep and sustained declines averaging 3-4% per year both range-wide and within most of the states and regions they occupy [4, 10]. This equates to a loss of about 85% of the population over 50 years, one of the largest recorded declines among all widely-distributed passerine birds in the interior western United States [10, 11].

**Fig 1.**
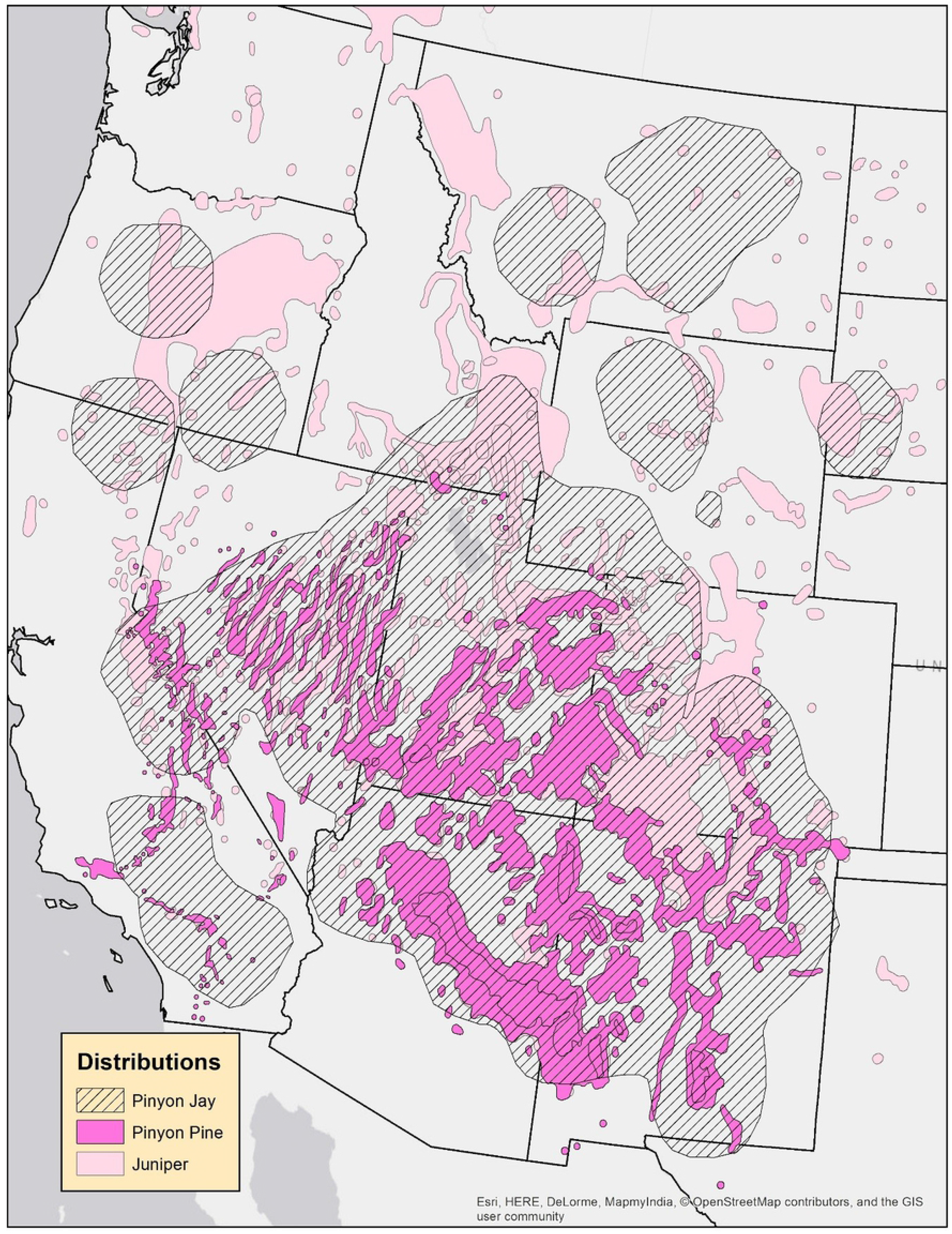
Distribution of the Pinyon Jay [4], pinyon pine (*Pinus monophylla* and *P. edulis* combined), and juniper (*Juniperus osteosperma*, *J. occidentalis*, and *J. spoculorum* combined) [5]. Some of the juniper distribution is invisible under the pinyon pine distribution on the map.

**Fig 2.**
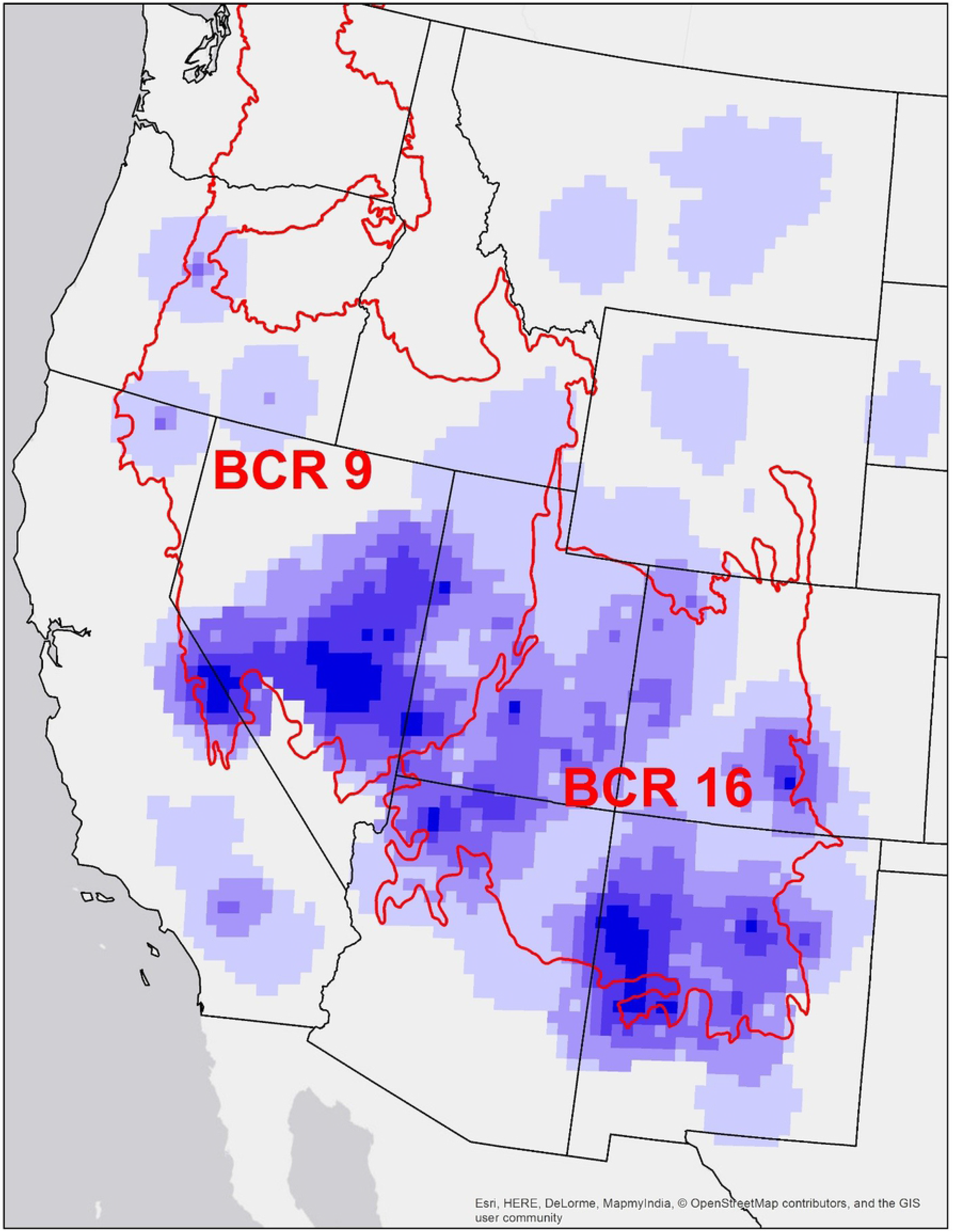
Relative density of the Pinyon Jay (purple colors, with darker colors representing higher relative densities [4]) and Bird Conservation Regions (BCRs) 9 (Great Basin) and 16 (Southern Rockies / Colorado Plateau).

Despite this decline, no systematic conservation efforts have been undertaken for the Pinyon Jay, although it is included on the ‘sensitive species’ lists of many federal and state management agencies and avian conservation organizations [12–15], and is the subject of a recent interagency working group conservation strategy [15]. The lack of conservation action for Pinyon Jays may be attributable to several factors. First, despite a rich knowledge of the species’ social behavior, breeding behavior, and spatial memory [6, 16, 17], its occupancy and habitat use patterns are poorly characterized and understudied in most parts of its range. Second, there are no widely-accepted or strongly-supported hypotheses about the causes of Pinyon Jay population declines. Finally, the landscapes inhabited by Pinyon Jays are primarily managed for other priorities. These include improving habitat for game species such as Greater Sage-Grouse (*Centrocercus urophasianus*) and mule deer (*Odocoileus heminus*), creating wildlife corridors, and mitigating fire hazards [3, 18–22].

Pinyon Jay declines could be related, at least in part, to changes in the pinyon-juniper woodlands that comprise most of their habitat [3, 22–24] and much of the forested landscape of the Great Basin (i.e., BCR 9) and Colorado Plateau (i.e., BCR 16) [25–28]. The spatial extent of pinyon-juniper woodlands (most commonly a *Pinus monophylla - Juniperus osteosperma* association) in this region has undergone climate-induced fluctuations since the end of the Pleistocene epoch 11,500 years ago [29, 30], but it has been suggested by some authors that over the last 150 years, extension of local woodland range (i.e. “expansion”) and increased tree densities within extant stands (i.e. “infill”) have occurred at atypical and perhaps unprecedented rates, at least in the Great Basin [22, 31–34]. Other authors have questioned this conclusion and suggest that expansion and infill are either localized, part of a historically normal pattern of spatio-temporal woodland dynamics, or represent recoveries from earlier widespread clearing during the western settlement period [22, 35, 36]. Regardless of which of these paradigms is correct, over recent decades extensive acreages of pinyon-juniper woodlands have been clear cut or thinned by resource management agencies and private owners to accomplish various management objectives [3, 37, 38].

In the Great Basin, the primary pinyon-juniper woodland management objective over the last 20 years has been creation or restoration of shrubland habitat by clear cutting stands that are regarded as encroaching into shrublands, often with a goal of benefitting Greater Sage-Grouse [39, 40]. If and how this type of vegetation management affects Pinyon Jays remains undetermined. Answering this question requires a better understanding of Pinyon Jay occupancy patterns and habitat use in the Great Basin, determining the extent of overlap between their preferred habitats and ongoing vegetation management activities, and monitoring or modeling the effects of those activities. In this study, we compared the habitats used by Pinyon Jays for caching or retrieving seeds, for foraging, and for nesting, to the full range of habitat available within pinyon-juniper woodlands. Our goals were to determine whether Pinyon Jays in the Great Basin used predictable subsets of available habitat for these distinct behavior types. If so, this information could assist managers seeking to incorporate Pinyon Jay conservation measures into existing vegetation management programs for pinyon-juniper woodlands, and help to guide future research [15].

## Methods

### Overview of study design and study region

We used a case-control study design [41, 42], with observed locations of Pinyon Jay flocks as case records and a set of pre-existing Forest Inventory Analysis (FIA) plots established by the U.S. Forest Service (USFS) as control sites (see below for details). Pinyon Jay data were collected from 2008 – 2014 in three Great Basin study areas. The control sites were distributed over a broader study region located entirely within the Central Basin and Range Ecoregion [43] and BCR 9 (Fig 2) that encompassed all three Pinyon Jay study areas (Fig 3). Control sites were not assumed to be unoccupied by Pinyon Jays, but were intended to provide a representative characterization of the diversity of pinyon-juniper woodland habitat available within the general study region. Habitat at Pinyon Jay locations and control sites was quantified using the standardized FIA protocol (see below) within circular 0.405-ha (1-acre) plots, which defined the spatial scale of this analysis. Analyses consisted of ordinations and logistic regressions, supplemented by data visualizations. We recognize the potential bias inherent in case-control sampling designs and followed Manly et al.’s [44] and Keating and Cherry’s [42] guidance for analysis.

**Fig 3.**
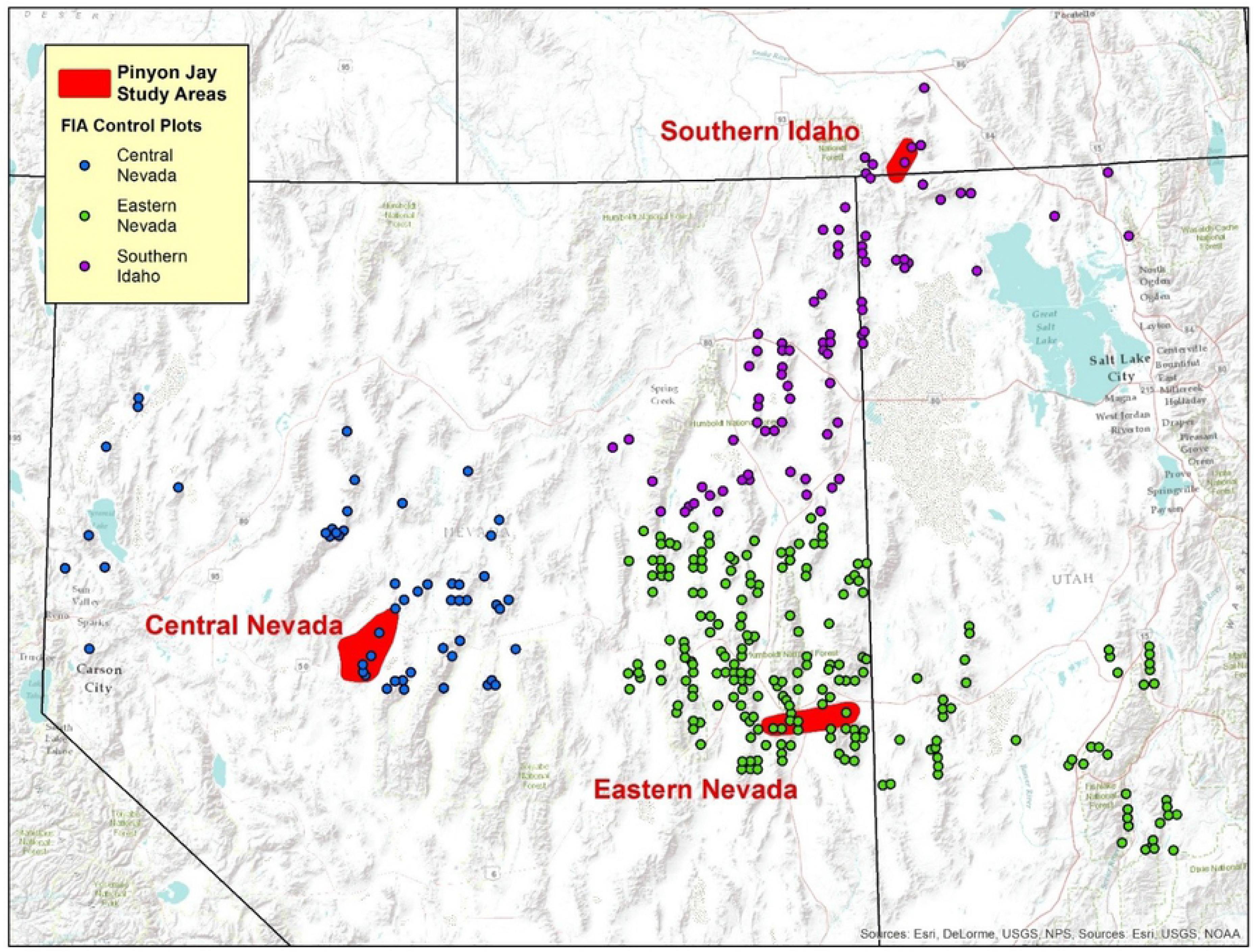
Locations of three Pinyon Jay study areas and FIA plots that served as control sites. FIA plot locations are approximate (i.e. “fuzzed”) to comply with USFS data protection policies. FIA plots are color coded to indicate the Pinyon Jay study area with which they were analytically paired.

### Pinyon Jay study areas and timeline

Pinyon Jay locations came from multiple Pinyon Jay flocks in these three study areas (Fig 3):

1. <u>Eastern Nevada (~ 63,130-ha study area)</u>, specifically the foothills between Baker, Nevada and Great Basin National Park, and Steptoe Valley south of the town of Ely, Nevada. This area is mostly comprised of public lands managed by the U.S. Bureau of Land Management, but includes some private property near Baker. Pinyon Jay data were collected from 2/28/2008 – 6/26/2008 and from 5/10/2009 – 8/27/2009.
2. <u>Southern Idaho (~ 23,470-ha study area)</u>, specifically the area in and around City of Rocks National Reserve and Castle Rocks State Park in Cassia County. This study area contains the northernmost occurrence of pinyon pine in North America [45]. Jurisdictions within the study area are the U.S. National Park Service, Idaho State Parks, U.S. Bureau of Land Management, and private lands. Pinyon Jay data were collected from 7/20/2012 – 10/5/2012.
3. <u>Central Nevada (~ 89,030-ha study area)</u>, specifically the Desatoya Range which lies along the border between Churchill County on the west and Lander County on east. This study area is comprised almost exclusively of U.S. Bureau of Land Management lands, with limited private inholdings. Pinyon Jay data were collected from 3/29/2013 – 6/5/2013.

Study area locations and periods of data collection were governed by three different funding agreements, and therefore seasonality was not standardized across all three areas. All study areas are characterized by mountainous “basin and range” topography, with plant communities dominated by big sagebrush (*Artemisia tridentata*) at lower elevations, pinyon-juniper woodlands at mid-elevations, and various mountain shrub and forest types at high elevations. Pinyon-juniper woodlands range in elevation from ~ 1,500 m – 2,600 m across all study areas, and are usually comprised of varying proportions of *P. monophylla* and *J. osteosperma*. Public lands in all study areas are managed for multiple uses and experience varying levels of livestock grazing and off-road vehicle travel.

### Pinyon Jay data collection and processing

Pinyon Jay data collection had several distinct components; initial searches, observational surveys, capture and radio-tagging, and radio-telemetry surveys. The observational surveys and radio-telemetry surveys generated the Pinyon Jay locations that are the basis of our analyses.

Initial searches of potential Pinyon Jay habitat (defined as pinyon-juniper woodlands and visually adjacent shrublands) were conducted during the first 1-2 weeks of field work at a given study area on foot and by vehicle to identify all or most of the Pinyon Jay flocks present. Because Pinyon Jay flocks are visually apparent from long distances, “noisy” (except when at the nest), and spatially segregated from one another, we regarded this a feasible goal. Search patterns used during initial searches were not systematized but were instead tailored to take advantage of local topography and access points. Areas searched were delineated on imagery maps to facilitate thorough coverage of pinyon-juniper woodland across a given study area. Upon detecting a flock, the observer usually maintained contact over a period of 1-3 h on each of several visits to obtain a preliminary and approximate delineation of the flock’s movement patterns and primary activity areas.

Flocks that were consistently detected during initial searches were then subjected to more intensive study by observational surveys and/or radio-telemetry surveys to obtain a sample of occupied locations. Observational surveys involved establishing visual contact with a flock; observing the flock with binoculars from a distance sufficient to prevent alteration of flock behavior (typically > 75 m); and recording approximately once per hour the point coordinates of the estimated centroid of the flock’s location along with estimated flock size and predominate behavior type (Table 1) whenever possible. The goal of observational surveys was to obtain locations across an entire daylight cycle at least once per week (usually assembled from several observation sessions conducted during different time periods on different days) for each flock over the duration of the data collection period. Coordinates of flocks were usually obtained by recording observer position with a GPS unit; recording a bearing to the visually-estimated flock centroid with a compass; measuring distance to the flock centroid by rangefinder or estimating distance by eye (for shorter distances < 25 m); and then plotting the flock’s estimated point location in GIS based on these parameters. In some cases, coordinates could be obtained directly by GPS after a flock vacated a previously-occupied location. As observers became increasingly familiar with the daily movement patterns of a flock, which tended to be consistent, their efforts were increasingly focused on the portions of the daily activity cycle that were more difficult to characterize.

**Table 1.** Description of Pinyon Jay behavior types that were recorded during observational and radio-telemetry surveys whenever possible.

| Behavior Type | Description |
| --- | --- |
| Caching | Birds observed either caching or retrieving previously-cached pine nuts or other similar food items. |
| Foraging | Birds observed collecting any type of new food item in trees, shrubs, or on the ground, including pinyon nuts, other plant material, or insects. |
| Nesting | Nest site or sites confirmed by direct observation or by confirmatory behaviors, such as carrying nest materials. |
| Roosting<sup>1</sup> | Birds present at a location during period(s) of darkness. |
| Loafing<sup>2</sup> | Birds observed resting, usually in trees, while not engaged in any other listed behaviors. |
| Flyover<sup>2</sup> | Entire flock flying in a directional manner, without landing. Short flight movements by one or a few birds while the flock was engaged in one of the other listed behaviors were not considered flyovers. |

<sup>1</sup>Due to small sample size, this behavior type was used only for ordination analyses.
<sup>2</sup>Observations for these behavior types were excluded from analysis.

Radio-telemetry surveys were also used to obtain Pinyon Jay locations for some of the studied flocks. Two methods were used to capture Pinyon Jays for radio-tagging; baited walk-in traps and mist nets. Traps were used where Pinyon Jays could be consistently drawn (determined by game cameras) to a supplemental food station baited with shelled peanuts, sunflower seeds, and dried corn. In these areas, a home-made wood and mesh walk-in trap (0.6 x 1.2x 0.9 m) with an open door was placed at the site, and bait spread periodically inside the trap to habituate birds to entering the trap. When habituation was sufficient (determined by camera traps), capture attempts were made by rigging the door for remote manual release, observing activity at the trap from a blind ~ 25 m from the trap, and releasing the door with a pull cord when Pinyon Jays were inside the trap. Mist-nets were used in areas where walk-in traps were not viable, specially in locations routinely visited by Pinyon Jays where nets could be deployed without being easily detected by birds. Mist nets (60 mm mesh) were arrayed either singly, as doubles, or stacked, depending on the geometry of the woodland opening where they were erected. In some cases, call playback was also used to try to draw birds into the nets. Any Pinyon Jays captured by walk-in trap or mist net were weighed and aged; standard aluminum leg bands (U.S. Geological Survey) were attached; and radio transmitters (Advanced Telemetry Systems model A2450) were glued onto feather stubble clipped to ~ 0.3 cm above skin level in the interscapular area. No more than six individuals from any single flock were radio-tagged, since data were collected to characterize flocks rather than individual birds. Captured birds were handled and processed only by experienced individuals holding a U.S. Fish and Wildlife Service master banding permit. Radio-tagged birds were manually tracked using a handheld three-element Yagi antenna with an Advanced Telemetry Systems R410 or R100 receiver. The goal of radio-telemetry surveys was to collect hourly locations over one entire daylight cycle per week for each flock, either in a single long session or over several sessions of 3 – 5 hours on different days and at different times. Most often, telemetry fixes from one or more radio-tagged flock members were used to approach the flock and establish visual contact, at which point locational and behavior type attributes were collected exactly as described above for observational surveys. Occasionally, flock locations were estimated by biangulation or triangulation of telemetry bearings that were post-processed using LOAS software (Ecological Software Solutions, LLC). In cases where the flock was not directly observed, behavior type was only recorded when it could be reasonably inferred based on previous observations (i.e. return to a nest colony location) or context (roosting locations at after sunset).

For a given flock, locations could be obtained by either observational surveys, radio-telemetry surveys, or both, but because both approaches ultimately relied on the same observational process, we regarded the resulting data as comparable. Initially, radio-telemetry was the preferred survey approach, but by later data collection periods we determined that visual contact with most flocks could routinely be established without reliance on radio-tagged flock members, and our efforts to capture birds for radio-telemetry were discontinued.

The full set of Pinyon Jay locations recorded during field work were filtered and processed prior to analysis. First, any locations where no behavior type was recorded were deleted. Next, all “flyover” locations and loafing locations (Table 1) were deleted, based on the premise that these could occur throughout and sometimes beyond the flock’s core home range. Finally, locations that represented the same flock engaged in the same behavior at the same location were combined as follows:

1. All nesting locations for a single flock were spatially averaged using the Mean Center tool in ArcMap 10.5 (ESRI, Redlands CA), based on the premise that at a given time, a flock has only one nesting area.
2. If caching, foraging, or roosting locations were closer together than 71.8 m, they were spatially averaged to generate a single location, as above. This was the smallest threshold distance sufficient to prevent any overlap between the 0.405-ha habitat plots used to assess habitat (see below), so this process smoothed Pinyon Jay locations to a spatial scale that matched the habitat assessment scale.

After these steps, 152 Pinyon Jay locations with recorded behaviors were retained for analysis (see Table S1).

Data collection procedures described in this section were primarily observational and were not submitted to or approved by an Institutional Animal Care and Use Committee. Field work that involved capture, radio-tagging, and release of Pinyon Jays was conducted under U.S. Department of the Interior Federal Bird Banding Permit # 22912, Idaho Fish and Game Department Collection Permit # 120724, and Nevada Department of Wildlife Collecting Permit # 29948.

### Control site selection

Control sites were selected from the pre-existing FIA data set of plots visited and measured between 2005-2013. FIA data provide a probabilistic and geographically unbiased assessment of forest / woodland attributes over time and space [46, 47] and a robust dataset to describe habitat for multiple species [48–52]. One FIA plot is randomly positioned within every cell of a sampling grid that covers all public and private forest land in the United States, with an overall density of one plot per 2,428 ha (6,000 acres) within the extent of that sampling frame [53]. Each year, 10% of these plots are surveyed or re-surveyed in the western U.S. using a spatially-interpenetrating sampling design that avoids conflation of spatial with temporal trends [46]. The criteria used to select the subset of FIA plots that were appropriate control sites for this study were as follows:

1. The FIA plot had to be < 200 km from the nearest Pinyon Jay location retained for analysis.
2. Presence of pinyon-juniper woodland within the FIA plot had to be confirmed by direct observation of the field assessment crew during the most recent assessment visit. A woodland type classification based solely on remote-sensing data was not sufficient to meet this criterion.
3. The site had to be located in both the Central Great Basin Ecoregion [43] and in BCR 9.
4. The most recent assessment visit had to have occurred in 2005 or later.

In total, 346 FIA plots from the larger data set met these criteria (Fig 3) and were included in our analysis (see Table S1). Each control site was assigned a regional attribute that corresponded to the closest Pinyon Jay study area (n = 212 for the Eastern Nevada region, n = 81 for the Southern Idaho region, and n = 53 for the Central Nevada region). Control sites were not distributed symmetrically around the Pinyon Jay study areas (especially for Southern Idaho) because their extent was constrained by the distribution of pinyon-juniper woodland and by the USFS site selection process.

### Habitat assessment

Forested habitat at Pinyon Jay locations was characterized using current FIA field protocols [54]. Site attribute measurements at Pinyon Jay locations on non-forested land followed procedures outlined for a FIA All-Conditions Inventory [55]. For all Pinyon Jay activity sites, FIA plot centers were placed at the Pinyon Jay location point coordinates and assessments were performed by USFS crews fully trained in FIA protocols and procedures. Control sites were assessed as part of routine USFS operations in various years between 2005-2013 [56–58].

FIA plot layout is based on a 0.405 ha circle that defines four subplots of 7.3 m diameter within which actual measurements are made. One subplot is located at the center of the larger circle, and the other three subplots have their centers equally spaced along its circumference (Fig 4). On a standard forested FIA plot, over 120 attributes are measured within each subplot to characterize location, condition, and vegetation [59]. Subplot data are then averaged or summed over subplots as appropriate and extrapolated to generate data at the whole-plot scale. The subset of FIA attributes that were considered for use in this analysis are presented and briefly described in Table 2.

**Fig 4.**
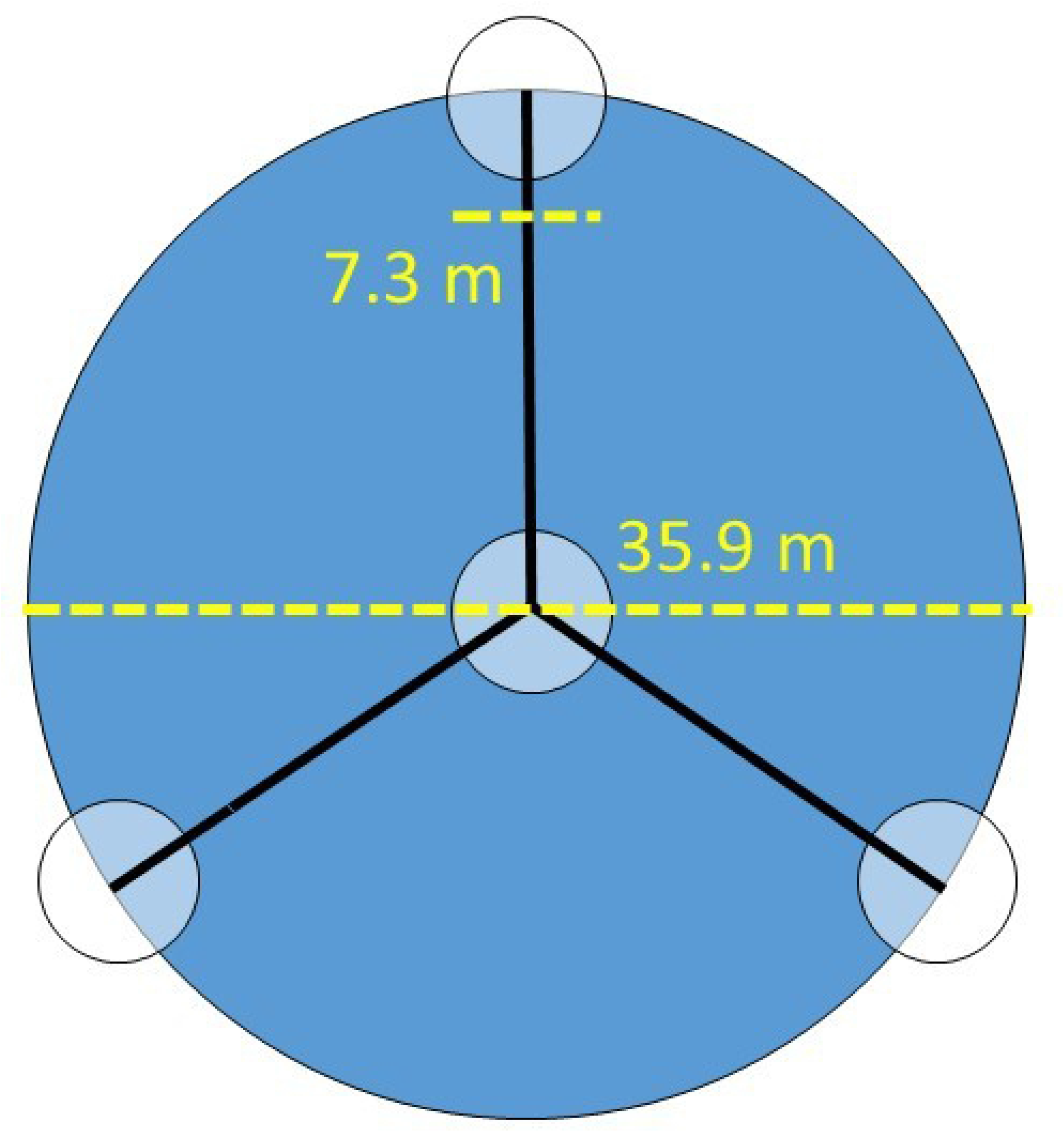
Schematic diagram of a USFS FIA plot layout. Data used in this study consisted of full-plot values produced by either summing or averaging sub-plot data.

**Table 2.** Attributes describing Pinyon Jay habitat that were considered in this study. Brief descriptions of attributes are provided parenthetically, with complete descriptions of associated methodologies available in [59]. The 2^nd^, 3^rd^, and 4^th^ columns indicate whether or not an attribute was used (Y = yes, N = No, NA = categorical attribute, not applicable) for the ordinations, logistic regressions, and data visualizations presented below. Footnotes provide additional explanations about attribute use considerations.

| Attribute | Ordination | Logistic Regression | Data Visualization |
| --- | --- | --- | --- |
| **FIA Attributes** |  |  |  |
| Elevation (measured in feet at plot center) | Y | Y | Y |
| Slope (expressed as slope percentage, or {{rise/run} x 100} at plot center) | Y | Y | Y |
| Habitat Type [60] | NA | N<sup>1</sup> | N |
| Stand Age (age in years of the oldest pinyon pine or juniper tree on plot, as determined by coring) | Y | Y | Y |
| Stand Density Index (index of three-dimensional tree density within stand [60]) | N<sup>2</sup> | N<sup>2</sup> | Y<sup>2</sup> |
| Canopy Cover (% by ocular estimation) | Y | N<sup>3</sup> | N |
| Tree Cover (% by ocular estimation) | Y | Y | Y |
| Shrub Cover (% by ocular estimation) | Y | Y | Y |
| Forb Cover (% by ocular estimation) | Y | Y | Y |
| Grass Cover (% by ocular estimation) | Y | Y | Y |
| Woody Debris (count of pieces of woody debris material along transects for seven different size diameter classes, defined by twig / branch diameter ranges) | Y | Y<sup>4</sup> | Y<sup>4</sup> |
| Distance to Road (km from plot center, assigned using topographic maps; original ordinal ranges converted to range midpoints) | Y | Y | Y |
| Disturbance Presence (“yes-no”, for any disturbances within the last five years) | NA | Y | N |
| Disturbance Type (categories of silvicultural treatment or other disturbance occurring within the previous five years) | NA | N<sup>5</sup> | N |
| **Non-FIA Attribute** |  |  |  |
| Distance to Edge (Distance to the lower-elevation woodland-shrubland ecotone, measured directly in on imagery in ArcMap 10.5) | N<sup>6</sup> | N<sup>6</sup> | Y |

<sup>1</sup> All sites used in analysis were characterized by presence of at least some pinon pine and/or juniper trees, but the FIA habitat type distinctions were too fine-grained (33 different types, median number of sites / type = 5) for inclusion in analysis.
<sup>2</sup>Omitted from analyses due to high correlations with other attributes, but included in data visualizations because of possible synthetic interpretational value.
<sup>3</sup>Omitted because of high correlation to Tree Cover attribute ( $r = 0.68$ ).
<sup>4</sup>Reduced to a single attribute by Principle Component Analysis that described 86.1% of variation across all original classes.
<sup>5</sup>The level of articulation was too fine for the limited number ( $n=68$ ) of disturbed sites and too unevenly distributed among types, so only the “yes-no” disturbance attribute was retained.
<sup>6</sup>Omitted from ordinations and logistic regression because attribute estimation process was not part of the standardized FIA protocol. Retained in data visualizations for possible interpretive value

Based on our field observations of Pinyon Jay landscape use over the course of the study, we decide to create one additional non-FIA attribute, “Distance to Edge” (Table 2) which is the shortest linear distance between a Pinyon Jay location or control site and the lower-elevation woodland-shrubland ecotone. First, polylines were digitized in ArcMap 10.5 to delineate the approximate ecotonal boundary based on visual examination of imagery. This process did not delineate woodland – shrubland boundaries inside well-defined woodland patches, but instead defined distinct woodland patches based on their lowest-elevation ecotonal boundary. Then, the shortest distance from each Pinyon Jay location or control site to the polyline was computed using the Near tool in ArcMap 10.5. This value was usually positive, but could be negative if a Pinyon Jay location was in shrubland at a lower elevation than the polyline. Because the Distance to Edge metric was not a FIA attribute, it was used only for data visualizations, not for statistical analysis.

### Analysis

#### Data

The complete data set used for all analyses and summarizations is provided in Table S1.

#### Ordination

To visualize how the habitat characteristics of Pinyon Jay locations for each behavior type overlapped with control sites, we performed region-specific ordinations on continuous FIA habitat attributes (Table 2) using nonmetric multidimensional scaling (NMDS). NMDS optimizes the non-parametric monotonic relationship among the data and the visualization space, providing the best low-dimensional representation [61–63]. All ordinated attributes were standardized with a z-transformation across all of the regions prior to the NMDS to promote optimization, which was defined as minimizing the stress value [63]. The NMDS was performed with the metaMDS function in the vegan package (v 2.5-5; [64]) in R (v3.5.1; [65]), using the Euclidian distance with two dimensions to force a visualizable representation. The maximum number of iterations was increased from the default of 200 to 10^10^ and the maximum number of runs was increased from 20 to 1,000 to achieve convergent solutions across repeated runs. Defaults for auto-transformation, extended dissimilarities, and weighted average scoring were turned off because they were not applicable to our transformed input data set.

#### Logistic Regression

To evaluate the effects of measured habitat attributes on Pinyon Jay occupancy by behavior type [42, 44, 66] we used logistic regressions for each behavior type that had sufficient data (i.e. caching, foraging, nesting). Roosting location were not analyzed because of low sample size (n=3). Because the ratio of Pinyon Jay locations to control sites was low, the odds ratio output from logistic regression can be treated as an approximation to the resource selection function [42, 67, 68]. However, it is important to recognize that the intercept value of the logistic regression does not estimate the overall use probability, as the Pinyon Jay locations were not randomly selected [42].

Initial evaluation of available predictor attributes (Table 2) consisted of plotting data pairs, evaluating correlations, and preliminary overall model fitting to ensure base-level convergence. Several attributes were eliminated from the analysis due to high correlations with other attributes or highly uneven distribution of values. Others were combined into a single attribute with principle components analysis (Table 2). All continuous attributes used for analysis (Table 2) were converted to metric units and z-transformed to facilitate model fitting and term comparison.

For the logistic regression models, we split the control sites assigned to each region (see above) according to behavior types in proportion to behavior frequency, rather than “reusing” control sites across multiple behavior types. Splitting was randomized and permutated 10,000 times to account for variation in the dataset splitting process. For each permutation, the control data were first split among the behaviors types within each region, then the data for a given behavior type were combined across the three regions and analyzed using a generalized (binomial) mixed model fit with the glmer function in the lme4 package (v1.1-19; [69]) in R (v3.5.1; [65]). The permutations were then combined to integrate over the sample allocation variation and estimate overall terms. This was done by first discarding any permutations that did not converge due to unreasonable splits of control sites that could occur within the randomization. Then, for each permutation, a single set of fixed effects parameters were drawn from the multivariate normal distribution described by the glmer model fit using the rmvnorm function in the mvtnorm package (v1.0-8; [70]) in R (v3.5.1; [65]). By drawing from the distribution within each permutation, the full set of parameter values across permutations includes both parameter estimation uncertainty and data allocation uncertainty. To avoid distributional assumptions in evaluating the significance of the parameter estimates, we tested whether the estimate significantly differed from 0 by using a permutation-style two-tailed approach [71]. This involved using the empirical cumulative distribution function (calculated using the ecdf function in the stats package in base R (v3.5.1; [65]) to estimate the two tail probabilities with respect to 0, taking the smaller value as the focal tail, and doubling that tail’s probability. We combined the estimates of the random effects (one for each successful permutation) across all of the permutations to generate the distribution of values for each random effects term across the uncertainty in control data allocation.

Within-sample classification accuracy of logistic regression models for each Pinyon Jay behavior type was averaged over all 10,000 permutations. Because the number of Pinyon Jay locations for each behavior type was limited, we did not withhold a subset of locations for external validation.

#### Data visualization

To aid in the interpreting statistical results and to highlight univariate patterns of interest, box plots were created for each continuous habitat attribute shown in Table 2, contrasting the distribution of attribute values for behavior-specific Pinyon Jay locations and control points.

## Results

### Data summary

Behavior-specific Pinyon Jay locations (n = 152) were obtained from 15 different flocks. Details about distribution of Pinyon Jay behavior type locations among study regions, apportionment of control sites, number of flocks per study area, and methods of data collection used within study areas are summarized in Table 3. Pinyon Jay data were not fully symmetrical among the study areas. Nesting was not recorded in Southern Idaho because the data collection period excluded the breeding season, and our data recording protocol in 2008-2009 for Eastern Nevada did not specify the foraging behavior type. Only caching locations were recorded in all three study areas, and only in the Central Nevada study area were all three major behavior types (caching, foraging, and nesting) recorded.

**Table 3.** Summary of Pinyon Jay locations after data processing, by behavior type and by region / study area. Shown parenthetically in 2^nd^ -5^th^ columns are the number of control sites assigned to each unique combination of study region x behavior type within logistic regression models as described in above. The final column summarizes the number of different Pinyon Jay flocks from which data were collected in each study area, with the data collection methods used shown in brackets (T = telemetry surveys, with number of deployed radio tags in parentheses; O = observational surveys).

| Study Region / Area | # Caching Sites | # Foraging Sites | # Nesting Sites | # Roosting Sites | # Flocks [Method of Study] |
| --- | --- | --- | --- | --- | --- |
| Eastern Nevada | 12 (106) | 0 (0) | 12 (106) | 3 (N/A) | 2 flocks [Flock #1 = T(6); Flock #2 = T(6)] |
| Southern Idaho | 32 (51) | 19 (30) | 0 (0) | 0 (N/A) | 2 flocks [Flock #1 = O & T(6); Flock #2 = O & T(2)] |
| Central Nevada | 26 (18) | 22 (15) | 28 (20) | 0 (N/A) | 11 flocks [All = O] |

### Ordination

The NMDS ordination found convergent solutions within 28, 388, and 20 runs for the Eastern Nevada, Southern Idaho, and Central Nevada regions, respectively (Fig 5). Each of the ordinations was well-correlated with true distances with stress values of 0.183, 0.127, and 0.161, respectively. Notably, for all three regions, Pinyon Jay locations formed distinct clusters in the NMDS plots that were easily distinguishable from the broader distribution of control sites (Fig 5) Degree of overlap among Pinyon Jay behavior type clusters varied somewhat across study regions, though nesting locations and caching locations were distinct in the two regions where both types of data were available. NMDS results suggested that the locations where Pinyon Jays occurred in the three study areas tended to be different than typical control sites within the corresponding study regions, and that Pinyon Jays tended to use different habitats for different behaviors.

**Fig 5.**
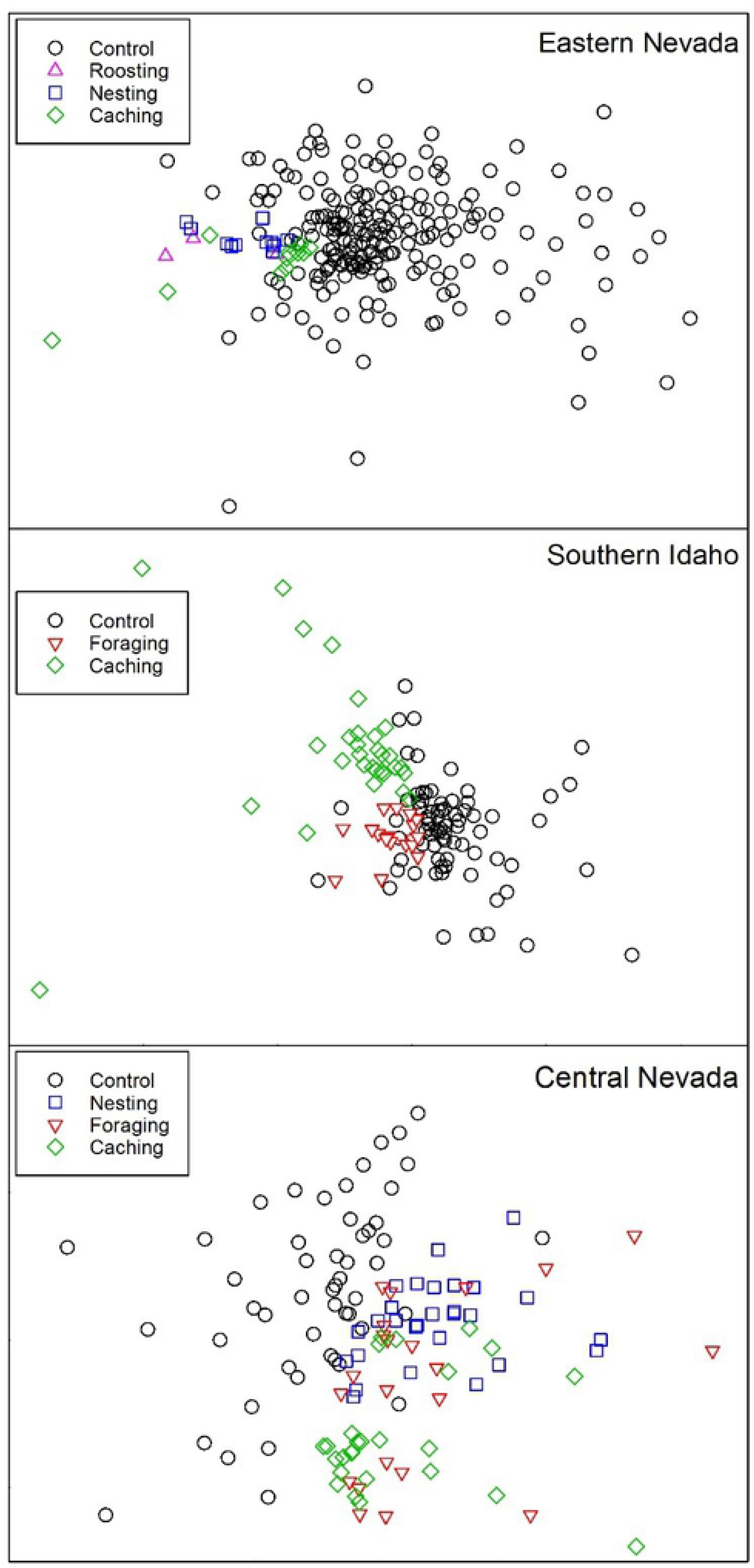
NMDS ordination results for each region, by location type. Note that each region’s ordination is independent and the axes cannot be compared among them.

### Logistic regression

Logistic regression models offer potential insights into which habitat covariates underpin the differentiation seen in the ordinations. We note, however, that within our logistic regressions, many predictors that had strong and statistically significant effects for a majority of iterations in the control site allocation process became non-significant when all iterations were combined. Because of this sensitivity to control site allocation, examination of data visualizations (next section) may assist in the interpretation of model results. Internal classification accuracy of behavior-specific Pinyon Jay locations versus control sites (averaged overall all iterations) was high (0.890 for the caching model, 0.863 for the foraging model, and 0.920 for the nesting model).

Models indicated that Pinyon Jay caching locations were more likely to occur with lower slope, lower tree cover, increased woody debris, shorter distance to roads, and more disturbance than control sites (Table 4). There was also a strong effect of decreased elevation, though it was not significant when averaged across all control allocation iterations. None of the predictors for Pinyon Jay foraging locations were statistically significant when averaged across all iterations due to high standard errors, but the largest effect sizes were noted for lower slope, less grass and forb cover, increased woody debris, and shorter distance to roads compared with control sites (Table 4). Pinyon Jay nesting locations were more likely to occur at lower elevations with decreased forb cover and increased woody debris compared to control sites (Table 4). Across all three Pinyon Jay behavior types, occupancy probability generally increased with lower elevation, lower slope, lower forb cover, shorter distance to roads, and increased woody debris compared to control sites (Table 4). Lower tree cover was a marginally significant predictor for caching locations but was not a significant predictor for foraging and nesting locations.

**Table 4.** Logistic regression results over all control site allocation iterations for the caching, foraging, and nesting location analyses. Fixed effects are reported as mean and standard error of the estimate and the two-tailed p value; random effects are reported as mean and standard error of the estimate and the probability (fraction of permutations) that the term was equal to 0. Bolded terms are significant at α = 0.05.

| **CACHING LOCATIONS** |  |  |  |
| --- | --- | --- | --- |
|  | Mean | Standard Error | p |
| Fixed Effect |  |  |  |
| **Intercept** | **-3.687** | **1.162** | **0.001** |
| Elevation | -0.405 | 0.585 | 0.481 |
| Slope | **-2.066** | **0.557** | **< 0.0001** |
| Stand Age | -0.177 | 0.432 | 0.654 |
| **Tree Cover** | **-0.913** | **0.553** | **0.075** |
| Shrub Cover | 0.251 | 0.349 | 0.445 |
| Forb Cover | -0.021 | 0.3 | 0.904 |
| Grass Cover | 0.377 | 0.439 | 0.385 |
| Woody Debris | **1.235** | **0.818** | **0.010** |
| **Distance to Road** | **-2.712** | **0.789** | **< 0.0001** |
| Disturbance | **1.971** | **0.951** | **0.022** |
| Random Effect |  |  |  |
| Eastern Nevada: Intercept | 1.838 | 0.413 | 0.0001 |
| Southern Idaho: Intercept | -1.320 | 0.301 | 0.0001 |
| Central Nevada: Intercept | -0.522 | 0.288 | 0.0001 |
| **FORAGING LOCATIONS** |  |  |  |
| Fixed Effect |  |  |  |
| Intercept | -5.486 | 7.418 | 0.044 |
| Elevation | -1.566 | 2.790 | 0.317 |
| Slope | -4.184 | 6.299 | 0.058 |
| Stand Age | -2.013 | 3.641 | 0.236 |
| Tree Cover | 1.931 | 3.079 | 0.200 |
| Shrub Cover | 1.380 | 2.479 | 0.311 |
| Forb Cover | -9.236 | 14.819 | 0.080 |
| Grass Cover | -4.970 | 8.384 | 0.113 |
| Wood Debris | 4.419 | 9.650 | 0.053 |
| Distance to Road | -3.326 | 4.848 | 0.063 |
| Disturbance | -1.700 | 75.272 | 0.555 |
| Random Effect |  |  |  |
| Eastern Nevada: Intercept | 0.686 | 1.076 | 0.290 |
| Southern Idaho: Intercept | N/A | N/A | N/A |
| Central Nevada: Intercept | -0.707 | 1.327 | 0.290 |
| **NESTING LOCATIONS** |  |  |  |
| Fixed Effect |  |  |  |
| **Intercept** | **-4.723** | **2.623** | **0.021** |
| **Elevation** | **-2.445** | **1.600** | **0.004** |
| Slope | -1.249 | 1.090 | 0.154 |
| Stand Age | 0.528 | 1.016 | 0.564 |
| Tree Cover | 0.500 | 1.030 | 0.570 |
| Shrub Cover | 0.261 | 0.848 | 0.743 |
| Forb Cover | **-7.110** | **4.247** | **0.022** |
| Grass Cover | 0.123 | 1.256 | 0.983 |
| **Woody Debris** | **2.156** | **1.266** | **0.003** |
| Distance to Road | -1.118 | 1.291 | 0.289 |
| Disturbance | 2.606 | 2.406 | 0.115 |
| Random Effect |  |  |  |
| Eastern Nevada: Intercept | 2.116 | 0.903 | 0 |
| Southern Idaho: Intercept | -2.088 | 0.904 | 0 |
| Central Nevada: Intercept | NA | NA | NA |

The random effects components of the three models were all estimated to be non-zero, although the foraging model estimated a zero-value in 29% of allocation iterations, whereas the caching model estimated a zero-value in < 1% of iterations. The additional complexity introduced by a singularity/boundary condition (a zero-value random effect) within many foraging model iterations may have contributed to the variability (and thus reduced significance) seen in the parameter estimates.

### Data visualizations

Fig 6 shows box plots for all FIA attributes included in logistic regression models combined across all study regions, and Fig 7 shows box plots for two additional habitat attributes (the FIA Stand Density Index attribute and the non-FIA Distance to Edge attribute) that were not included in logistic regressions, but which showed patterns of interest with regard to Pinyon Jay occupancy. Notable patterns in these box plots are as follows:

1. Across all study regions, locations used by Pinyon Jays appeared to be a distinct subset available woodland habitat with regard to many habitat attributes. This is consistent with the patterns observed in the NMDS ordination (Fig 5). Additionally, locations used by Pinyon Jays for different behaviors appeared to vary with regard to multiple habitat attributes.
2. Compared to available habitat, caching locations were concentrated in lower elevation, lower-slope areas with young woodland stands, low tree cover, and high shrub, forb, and grass cover. The Stand Density Index was typically very low for caching locations, which were also highly concentrated near the woodland-shrubland ecotone (i.e. low Distance to Edge values). In fact, many caching locations occurred in the pure shrubland habitat located down-slope from the woodland-shrubland ecotone (note that these pure shrublands were not represented in the sample of FIA control sites).
3. Foraging locations were also concentrated in lower elevation and lower slope areas, but were somewhat higher and steeper than caching locations, with somewhat higher stand densities. Foraging locations had more woody debris than typical control sites, but stand age, tree cover, forb cover, and grass cover were comparable to control sites. Shrub cover in foraging locations was highly variable, but tended to be higher than in control sites. Foraging locations were close to the woodland-shrubland ecotone, with lower stand density indices, though to a less extent than caching locations
4. Nesting locations also tended to occur at lower elevations and lower slopes, but to a lesser degree than foraging and caching locations. Stand age and shrub cover were comparable between nesting locations and control sites, but nesting locations had higher tree cover, lower forb cover, and more woody debris. Stand density was somewhat lower for nesting locations than for control sites, but with considerable overlap. Nesting locations were also concentrated close to the woodland-shrubland ecotone, but to a lesser degree than foraging locations.
5. There was a distinct pattern of increasing elevation and slope, increasing distance from edge, and increasing stand density moving from caching locations to foraging locations to nesting locations, but all three behavior types had lower values for these attributes than typical control sites.

**Fig 6:**
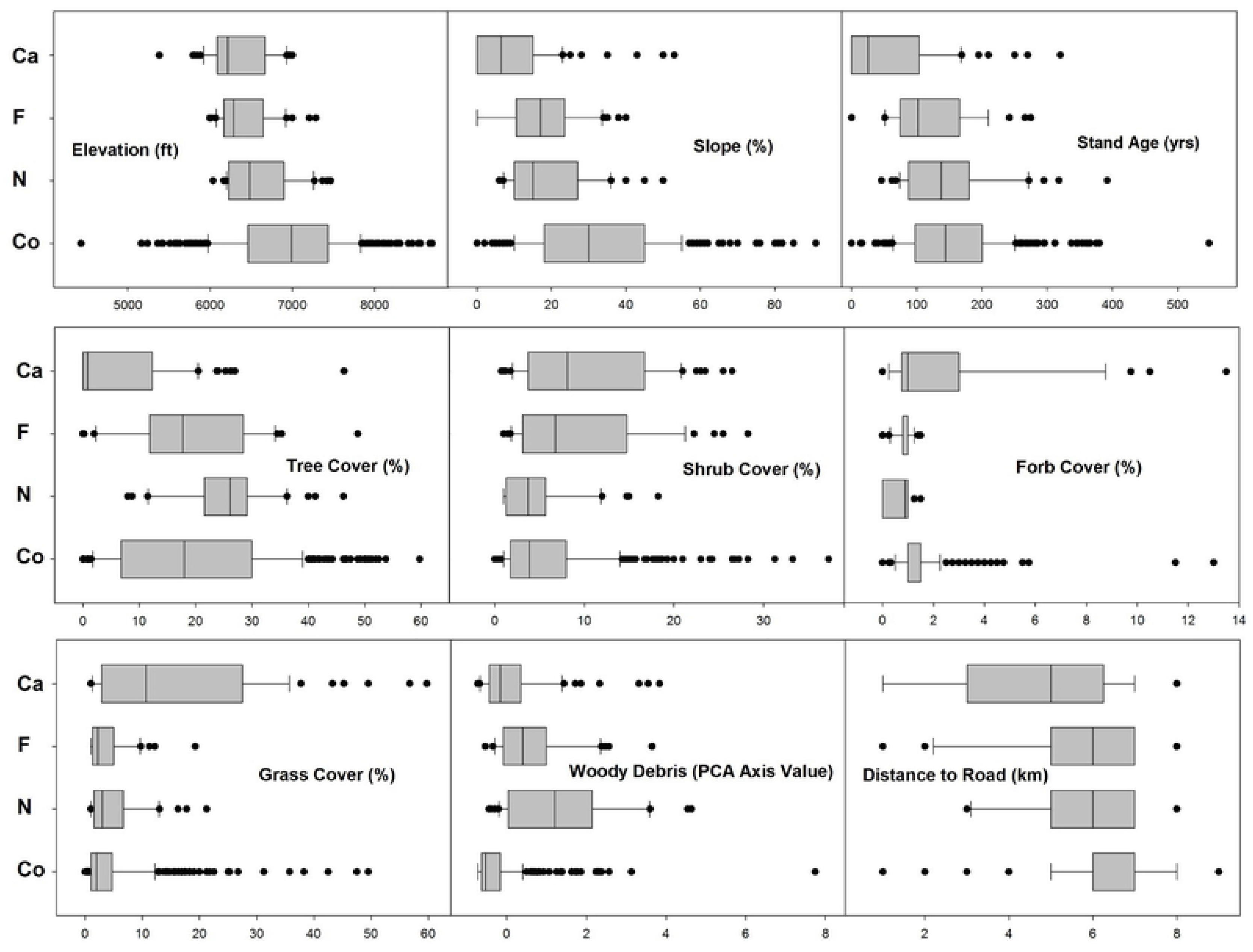
Box plots for all continuous attributes used in logistic regression models, as described in Table 2. Y-axis codes are Co = control sites, N = Pinyon Jay nesting locations, F = Pinyon Jay foraging locations, and Ca = Pinyon Jay caching locations. For better visual clarity, the extreme high range of observed Forb Cover values (15 – 25%) is truncated, omitting a small number of Co sites and Ca locations.

**Fig 7:**
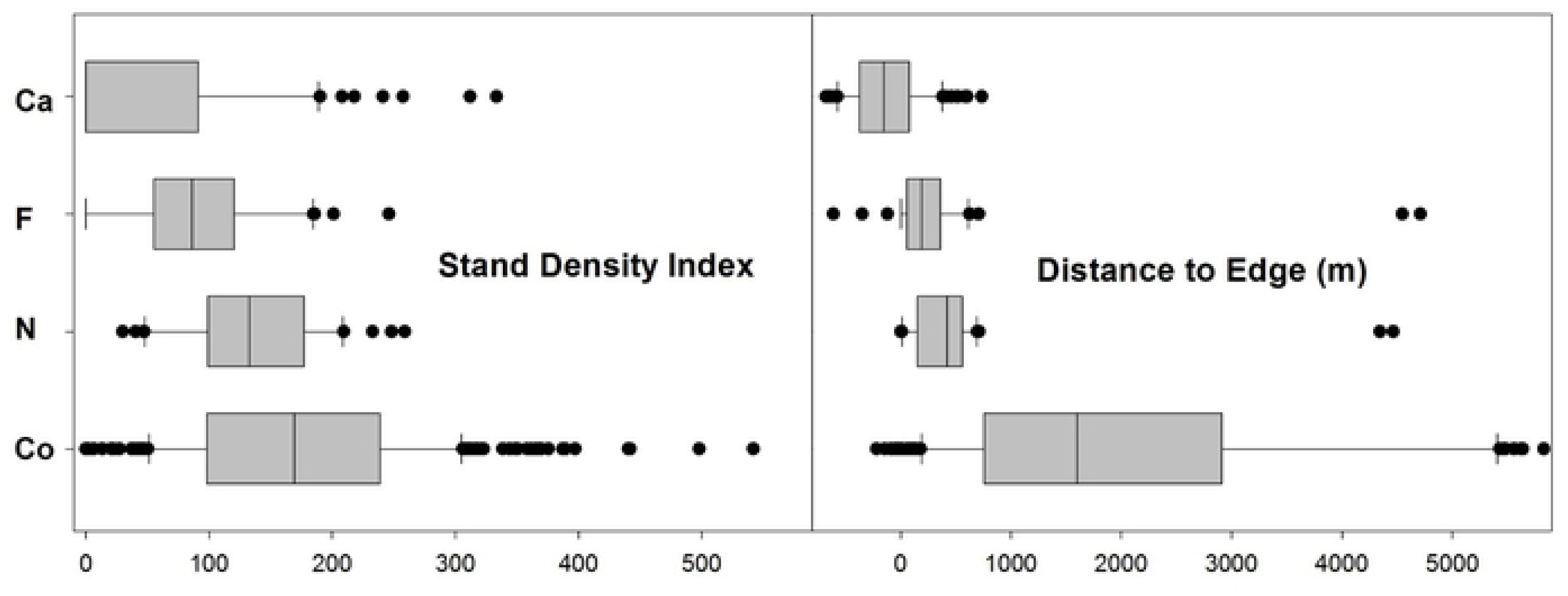
Box plots for two habitat attributes not used in logistic regression models, as described in Table 2. Y-axis codes are Co = control sites, N = Pinyon Jay nesting locations, F = Pinyon Jay foraging locations, and Ca = Pinyon Jay caching locations. For better visual clarity, the extreme high range of observed Distance to Edge values (6,000 – 12,000 m) values are truncated, omitting a small number of Co sites.

## Discussion

### Main findings and significance

This study offers the first systematic description of Pinyon Jay occupancy patterns and behavior-specific habitat characteristics in the Great Basin. The combination of ordinations, logistic regressions, and data visualizations presented in this study suggest that Pinyon Jays use pinyon-juniper woodlands selectively. All Pinyon Jay locations in our study areas were concentrated in or near lower-elevation, flatter woodlands and were less common in higher-elevation, steeper woodlands. Additionally, the areas used for different Pinyon Jay behaviors appear to have distinctive (but overlapping) habitat profiles, with caching, foraging, and nesting arrayed sequentially along gradients of increasing slope, elevation, and stand density. Similar patterns of Pinyon Jay caching and nesting activities partitioned along elevation and stocking gradients were observed by Johnson et al. [23] within a pinyon-juniper (*P. edulis* / *Juniperus spp.* association) woodland system in New Mexico, suggesting that habitat partitioning by behavior along an elevational gradient could be present across a broader physiographic range than our study regions.

Pinyon Jay caching locations were concentrated in open woodland stands with high shrub and grass cover, which are similar to the Phase I (early successional) pinyon-juniper woodlands defined in the classification scheme by Miller et al. [33], and sometimes occurred in pure shrublands. This occurrence pattern could have a mutualistic explanation [3]. From the pinyon pine perspective, seedlings likely experience less competition from established trees in more open areas, and seeds placed next to shrubs, rocks, or woody debris in otherwise open areas may benefit from favorable microsite conditions created and maintained by those features [72, 73]. From the Pinyon Jay perspective, pinyon pine seeds cached away from their source of origin may be less likely to be discovered and eaten by small mammals that specialize on pinyon seeds [73, 74].

Pinyon Jay foraging locations generally occurred in older (though still relatively young) stands than caching locations, across a wide range of tree cover values. Given that foraging behavior as defined in our study encompassed the gathering of diverse food items from trees, shrubs, and ground, areas with these characteristics may offer a beneficial combination of pinyon pines in their most productive seed-bearing years [75, 76] interspersed with areas where abundant insect prey is available due to higher shrub or ground cover [77]. The habitat characteristics of Pinyon Jay foraging locations correspond to a mosaic of Phase I and Phase II pinyon-juniper successional stages [33].

Pinyon Jay nesting locations tended to be concentrated in areas with higher tree cover and more woody debris, presumably because of the concealment they offer [23, 78]. However, like caching and foraging locations, nesting locations were concentrated in lower elevation, lower slope areas, and steeper, higher sites that otherwise offered good concealment for nesting locations appeared to be avoided. Pinyon Jay nesting locations correspond best with the denser portion of the Phase II class of pinyon-juniper woodlands, but may also include some Phase III areas [33].

All Pinyon Jay locations, regardless of behavior type, were concentrated in lower-elevation woodlands (most likely a mix of Phase I and Phase II classes) near the woodland-shrubland ecotone. This could occur because of the longer snow-free season of lower elevations, or the presence of a mosaic of desirable habitat characteristics needed to support different behavior types. Phase I and Phase II woodlands are relatively common at lower elevations where Pinyon Jay locations are concentrated, whereas the proportion of Phase III woodlands tends to increase with increasing elevations based on our field observations.

In addition to providing important information about Pinyon Jay occupancy patterns and habitat use, these findings are potentially significant for vegetation management planning and implementation because Pinyon Jays in our three study areas appear to prefer the same lower-elevation, relatively-open woodlands where most woodland removal management is performed [79, 80]. These vegetation management projects, which are most often conducted to create or improve habitat for Greater Sage-Grouse [81] (but see Miller et al. [22] and Somershoe et al. [15] for others reasons for woodland treatments), have resulted in the removal of an estimated 45,000 ha of pinyon-juniper woodland in the Great Basin portions of Nevada, Utah, and Idaho over the last eight years alone (Witt, unpublished USFS data). The pace of this activity appears to be steady or accelerating, and may therefore be or become a significant factor affecting Pinyon Jay populations, either negatively or positively. Information from other regions suggests that woodland treatments can have unintended effects on Pinyon Jays. For example, Johnson et al. [82] found that a fuels treatment within pinyon-juniper woodlands in northern New Mexico that reduced tree density by almost 90 percent prompted the local Pinyon Jay flock to avoid the treated area altogether. To date, however, almost no direct monitoring has been conducted in the Great Basin to determine if and how Pinyon Jay flocks respond to vegetation management projects that occur within or close to their home ranges.

### Interpretational considerations

The findings presented in this study should be interpreted with the following considerations in mind:

1. Pinyon Jay data were collected at the three distinct study areas that collectively encompassed 175,630 ha. Although this was a large area, it represents only a small portion of the Great Basin region that we would like to characterize.
2. This study combined data from three distinct projects that covered different years and seasons. This is most immediately relevant to interpreting the foraging model, given that seasonal variation in foraging behaviors and locations are plausible.
3. We equated “potential habitat” for Pinyon Jays to all pinyon-juniper woodlands lying within 200 km of any Pinyon Jay study area, as sampled by FIA plots. Although inferences from this study can cautiously be extended to the pinyon-juniper woodlands beyond our immediate Pinyon Jay study areas (subject to confirmation in future studies; see Johnson and Sadoti [78] for caveats about model transferability to similar systems) it does not extend to other forest types or regions where Pinyon Jays occur.
4. Our analysis does not distinguish between flocks within a study area, and does not allow us to draw inferences about inter-flock variability with regard to behavior-specific occupancy patterns.
5. The iterative allocation of control sites among Pinyon Jay behavior types in logistic regression modeling was necessary, but it could have diluted the statistical significance of some important predictors of occupancy. Future modeling efforts where the spatial extents and sample sizes of Pinyon Jay data and control data are better matched should reduce this issue. Models created from data sets with a large sample of Pinyon Jay locations will also allow data to be withheld from the model building process and used for external validation.
6. Interpreting the relationship between Pinyon Jay occurrence and distance to road is complex and potentially non-causal. Roads density is typically higher in lower-elevation, flatter areas, and Pinyon Jays prefer these areas for reasons completely unrelated to the proximity of roads. However, one author has suggested the vegetation typically present alongside graded, unpaved roads may provide valuable foraging opportunities for Pinyon Jays [83].
7. The Distance to Edge attribute used for data visualizations shows clear contrasts across behavior-specific Pinyon Jay locations and control sites, but it needs to be further investigated using more formal methods for delineating ecotones. Additionally, the patterns seen in our data (Fig 7) were enhanced because control sites by definition excluded the pure or near-pure shrublands that comprised a significant proportion of Pinyon Jay caching locations, and a smaller proportion of foraging locations.

Currently, we are analyzing a separate Pinyon Jay data set derived from a long-term statewide bird monitoring program in Nevada. Because these data were obtained from a broader-scale fully-randomized sampling design and used a fully standardized survey protocol, they should provide a useful independent characterization of Pinyon Jay occupancy patterns in the Great Basin.

### Conclusions and recommendations

Pinyon Jay populations have been declining precipitously for at least the last half-century, while the pinyon-juniper woodlands that they inhabit in the Great Basin are thought by many to have been expanding at unprecedented rates [84–87]. Given this apparent paradox, identifying the reasons for Pinyon Jay declines is critical for defining constructive conservation actions that ensure the species’ long-term viability [3, 15]. This urgency is amplified by the widespread and potentially accelerating woodland management activities in the Great Basin that prioritize creation or preservation of shrublands without a clear understanding of their impacts on Pinyon Jays and other woodland-associated birds. Progress towards a more inclusive management paradigm can be achieved through: a) better knowledge of Pinyon Jay ecology and habitat requirements, b) monitoring of management impacts, c) better understanding of the ecology and dynamics of the woodlands that comprise Pinyon Jay habitat, and d) integration of knowledge obtained from these three areas into existing vegetation management protocols and guidance.

A fundamental need is for more robust, spatially-extensive data characterizing Pinyon Jay occupancy patterns and habitat use as a function of region, season, and behavior type, both across and within the individual flock level. To maximize their value, these data would ideally be gathered using a standardized survey protocol. Pinyon Jays, however, present multiple challenges to the field biologist, study designer, and data analyst [15], and approaches suitable for typical passerine birds may be suboptimal for Pinyon Jays for several reasons. Unlike species where a single breeding pair occupies a clearly-delineated territory at all times during the breeding season, Pinyon Jays occupy (and presumably select) habitat at the flock and subflock level. Pinyon Jays are also year-round residents, and protecting breeding habitat alone may be insufficient for effective conservation. The Pinyon Jay’s pattern of habitat use, which involves temporal flock movements across a relatively large home range to accommodate different behaviors and take advantage of seasonally-varying food resources, has potentially profound effects on both the detection properties of a given survey protocol and the ecologically-legitimate interpretation of those detection patterns. As a simple example, a Pinyon Jay flock may be frequently absent from a critically-important subset of its home range, either during portions of the day, or during entire seasons. Similarly, roaming flocks may frequently fly over or loaf in areas of the home range that are not critical to home range quality or viability. To provide accurate and actionable information about the habitat requirements of Pinyon Jays, survey protocols and research study designs need to account for these realities appropriately, operate at scales that reflect actual Pinyon Jay habitat selection patterns, and be guided by a sampling framework that produces well-balanced data suitable for presence / absence modeling. The multi-agency Pinyon Jay Working Group [15] is currently actively investigating options for standardized Pinyon Jay survey protocols. The same group’s Pinyon Jay Conservation Strategy [15] also noted that some of the information needed to better characterize Pinyon Jay habitat requirements could be obtained by systematically monitoring Pinyon Jay responses to vegetation management activities, especially in situations where their pre-treatment presence has been confirmed by baseline or clearance surveys.

With regard to pinyon-juniper woodlands, their structural attributes and other characteristics that might be limiting to Pinyon Jay populations need to be further studied. We suggest that it may be especially important to identify the correlates or profiles of tree stands and landscapes that exhibit a predictable and/or abundant pinyon pine mast [22]. Our preliminary review of FIA data collected in Nevada between 2006-2015 suggests that woodlands matching the structural characteristics Pinyon Jays used for foraging in this study are 5-7 times less extensive than nesting or caching habitat in Nevada (unpublished USFS data). Presence of reliably productive stands within the home range could be especially important to Pinyon Jays during years of more generally depressed pine mast production. Given evidence of reduced mast production in pinyon pine [88], and associated changes in habitat use by Pinyon Jays [24] in some areas affected by climate change, it might be critical to long-term Pinyon Jay conservation to systematically investigate the quality and quantity of good foraging areas.

In addition, more research is needed to better clarify the degree to which woodland expansion and infill is part of a historically “normal” dynamic, versus a problematic departure condition. In the absence of this more holistic understanding, colonization of shrublands by trees tends to be widely regarded as “invasive”, even though at least some of these recently colonized areas appear to be an important component of Pinyon Jay habitat in the Great Basin. Achieving this broader perspective may be a necessary prerequisite to successfully accommodating the needs of Greater Sage-Grouse, Pinyon Jays, and other sensitive shrubland and woodland bird species within the overall framework of pinyon-juniper woodland management.

Ultimately, accruing information about Pinyon Jays must be incorporated into woodland management paradigms and protocols in the Great Basin (see Ricca et al. [89] for an example) in ways that accommodate both previously-identified and newly-emerging goals within the context of healthy ecosystem function [90]. For the present, Somershoe et al. [15] provide guidance for managers seeking to incorporate Pinyon Jay conservation measures into their vegetation management projects based the extent of current knowledge.

## Supporting Information Captions

S1 Table. All data used for data visualizations, ordinations, and logistic regression analysis, as summarized in Table 2. Each record refers to either a behavior-specific Pinyon Jay location or a control site. Each record includes various locational and data category attributes, including decimal latitude and longitude, along with all habitat attributes considered for inclusion in analysis, as described in Table 2. All attribute headers are sufficiently explicit to be self-explanatory when viewed in conjunction with Table 2. Definitions of codes for habitat types (which were not used in any analysis or data visualization) are available in Alexander 1988 [60]. Missing data are intrinsic to the FIA data set.

## Acknowledgments

Field work was conducted by Gustavo Gonzalez, Michael Maples, and John B. Free (in Eastern Nevada), by Murrelet Halterman, Natasha Peters, Larry Teske, and Mercer Owen (in Southern Idaho, and by Mercer Owen and Sue Brunner (in Central Nevada). Their efforts, which were supplemented by the authors, are much appreciated. Statistical analysis was performed by Juniper Simonas of Dapper Stats (www.dapperstats.com).

Wallace Keck, Superintendent of City of Rocks National Reserve and Park Manager for Castle Rock State Park, provided us with extensive information about local Pinyon Jay populations, generously allowed us use of park facilities, with further assistance from Trenton Durfee. Many local landowners permitted us to access to their property, notable among them LeAnn and Kim Draper, whose property functioned as a capture location and staging area for telemetry efforts. At the Bureau of Land Management, Sandra Brewer and John Wilson provided invaluable assistance and support in Central Nevada. We thank all of these individuals.

